# Genome-resolved ecology and evolution of the microbiome enabled by quantitative long-read metagenomics

**DOI:** 10.64898/2026.09.28.754546

**Authors:** M. Omar Din, Caitriona Brennan, Jianshu Zhao, Renee Oles, Lucas Patel, Amanda Birmingham, Dilara M. Sevilir, Lauren Hansen, Gillian M. Wright, Madison Ambre, Meika Tomita, Tara Boyer, Antonio González, Gail Ackermann, Daniel McDonald, Sarah Kralicek, Gail Hecht, Jeff Hasty, Larry Smarr, Rob Knight

**Affiliations:** Department of Pediatrics, University of California, San Diego; La Jolla, CA, USA; Synthetic Biology Institute, University of California San Diego, La Jolla, CA, USA; Division of Biological Sciences, University of California San Diego, La Jolla, California, USA; Department of Pathology, University of California, San Diego, La Jolla, California, USA; Bioinformatics and Systems Biology Program, University of California, San Diego; La Jolla, CA, USA; Medical Scientist Training Program, University of California, San Diego, La Jolla, CA, USA; Department of Biology, University of California, San Diego, La Jolla, California, USA; Department of Biomedical Sciences, Midwestern University, Downers Grove, IL, USA; Division of Gastroenterology, Department of Medicine, Loyola University Medical Center, Maywood, Illinois; Department of Microbiology and Immunology, Loyola University Chicago, Maywood, Illinois; Department of Bioengineering, University of California San Diego, La Jolla, CA, USA; Biodynamics Laboratory, University of California San Diego, La Jolla, CA, USA; Department of Computer Science and Engineering, University of California San Diego, La Jolla, CA, USA; Center for Microbiome Innovation, University of California San Diego, La Jolla, CA, USA; Halıcıoğlu Data Science Institute, University of California San Diego, La Jolla, CA, USA; Hong Kong University of Science and Technology Jockey Club Institute for Advanced Study, Hong Kong University of Science and Technology, Hong Kong SAR, China

## Abstract

Standard metagenomics resolves microbial communities only in relative terms and at coarse taxonomic resolution, obscuring how individual strains change in absolute abundance and evolve within a host^1–6^. We present long-read quantiomics (LRQ), a quantitative metagenomics platform coupling long-read sequencing with plasmid-based internal standards to recover high-quality genomes, including from degraded or low-input stool, and measure absolute, genome-resolved strain abundances. We validate LRQ using large plasmid standards and bacterial spike-ins, then apply it to eight years of dense sampling from an individual with colonic Crohn’s disease. LRQ resolves co-resident strains of the same species following opposing trajectories with inflammation. Escherichia coli, long treated as a single pathobiont, separates into two sub-populations with opposing inflammation responses; the inflammation-associated lineage carries an immune-evasion module of virulence-associated metal-acquisition, biofilm, capsule and toxin genes. We observe genome-wide adaptive sweeps on timescales matched to disease incidence. LRQ therefore converts long-read metagenomics from a genome-recovery tool into a quantitative framework providing absolute, strain-level measurement of microbial ecology and evolution.

## INTRODUCTION

In biology, mechanistic understanding requires knowing not only which components are present, but their absolute abundances and how they change over time. Copy number and concentration shape cellular behaviour, species interactions, and the dynamics of biological communities. In microbiome research this distinction is critical: relative abundance measurements can obscure underlying changes in microbial load ^7^. A taxon may appear to rise or fall simply because the abundance of other taxa has changed, confounding biological variation with compositional effects ^8^. Absolute quantification is therefore essential for interpreting community dynamics, comparing samples across conditions, and linking microbial change to host or environmental phenotype ^9^.

Most microbiome methods resolve communities only at broad taxonomic levels such as family, genus, or species ^1^. Yet functional and ecological properties often segregate below the species level, where closely related strains can differ in metabolism, virulence, and antimicrobial resistance ^1,10^. Existing techniques therefore risk grouping organisms with distinct, even opposing, effects on the host. Evolutionary dynamics are still less tractable, since they require resolving mutations and haplotypes within specific strains. Together these limitations motivate an approach that resolves absolute strain-level dynamics from high-quality genomes.

Previous absolute quantification protocols are mostly based on qPCR or sequencing of marker genes such as 16S rRNA or total microbial load measurements via flow cytometry ^7,11–14^, rather than quantification based on whole genomes. Targeted approaches can lead to inaccurate cell counts when marker genes are multi-copy, or are limited to high levels of taxonomic resolution. Shotgun metagenomics can enable large-scale surveys of taxonomic and functional diversity ^15^, but still yields relative abundance data rather than absolute quantification. Furthermore, short reads often fail to span repetitive or conserved regions, full genes, and operons, leading to fragmented and low quality assemblies ^16^ and ambiguous mapping ^17^, particularly among closely related species and strains ^1,18^. Whole genome sequences or cells can also be used as spike-in DNA standards ^19–21,22,23^, but this approach is not community-agnostic because samples may contain genomic sequences that resemble the spiked-in sequences and interfere with quantification accuracy. A plasmid-based approach with synthetic sequences can address this problem ^24^, although similar limitations exist due to short-read ambiguity and multi-mapping to the plasmid backbone or cloning strain DNA. Alternative strategies attempt to mitigate these challenges using entropy-based thresholds and mapping corrections using spiked-in synthetic DNA standards ^25^.

Long-read metagenomics is poised to dramatically expand our ability to accurately identify and assemble high-quality microbial genomes directly from complex samples ^2,26,27^. It also facilitates absolute quantification by accurate mapping of reads to genomes and quantification spike-in standard sequences. Although long-read metagenomics is rapidly emerging as a research focus, current approaches lack standardized, high-throughput protocols and absolute quantification methods that can be broadly applied across microbiomes. Here we develop long-read quantiomics (LRQ), a long-read metagenomics framework for absolute microbiome profiling that addresses existing limitations. LRQ combines long-read metagenomic sequencing with a synthetic DNA plasmid spike-in standard to enable genome-resolved quantification. Improved assembly and mapping properties of long reads also allow tracking of microbial pathogens, and generation of long haplotypes at SNP resolution. Applied to 8 years of dense longitudinal sampling from a patient with colonic Crohn’s disease, LRQ uncovers strain-level ecological and evolutionary dynamics—including co-resident lineages of Escherichia coli tracking inflammation in opposite directions, and adaptive sweeps that accelerate on the timescale of a flare—that are inaccessible to relative abundance alone.

## RESULTS

To make strain-level microbial behavior measurable at an absolute scale rather than only relative, we developed a quantitative long-read metagenomics approach using an automated preparation protocol for PacBio libraries that addresses common challenges with obtaining and processing high-molecular weight (HMW) for metagenomic samples. We found that the recently developed Matrix-tube extraction method generates high-quality DNA from stool, with >80% of fragments >10 kb in non-degraded samples (see Methods)^28^. Because metagenomic samples are frequently degraded, we developed a rapid, fully automated protocol to remove short DNA fragments in minutes, thereby rescuing samples for amplification and long-read sequencing (Fig. 1a, bottom left). This protocol is portable for metagenomics applications, such as marine and wastewater (see Methods) ^29,30^. To enable quantification, we use plasmids carrying unique, synthetic sequences, which can therefore be used as quantification standards for any microbiome (Fig. 1a, top left). We can accurately map the long reads to closely related metagenome-assembled genomes (MAGs) and quantification spike-ins, enabling accurate delineation of a standard curve and subsequent quantification of individual MAGs (Fig. 1b). This paradigm of quantitative, long-read metagenomics allows us to pursue investigations at the genome level, including strain-level dynamics and evolution (Fig. 1c). Here, we consider these assembled genomes as strain-level units which at most share 99.5% average nucleotide identity (ANI), a threshold that has been empirically validated to be most consistent with discrimination of intra-species diversity^31^.

**Figure 1:**
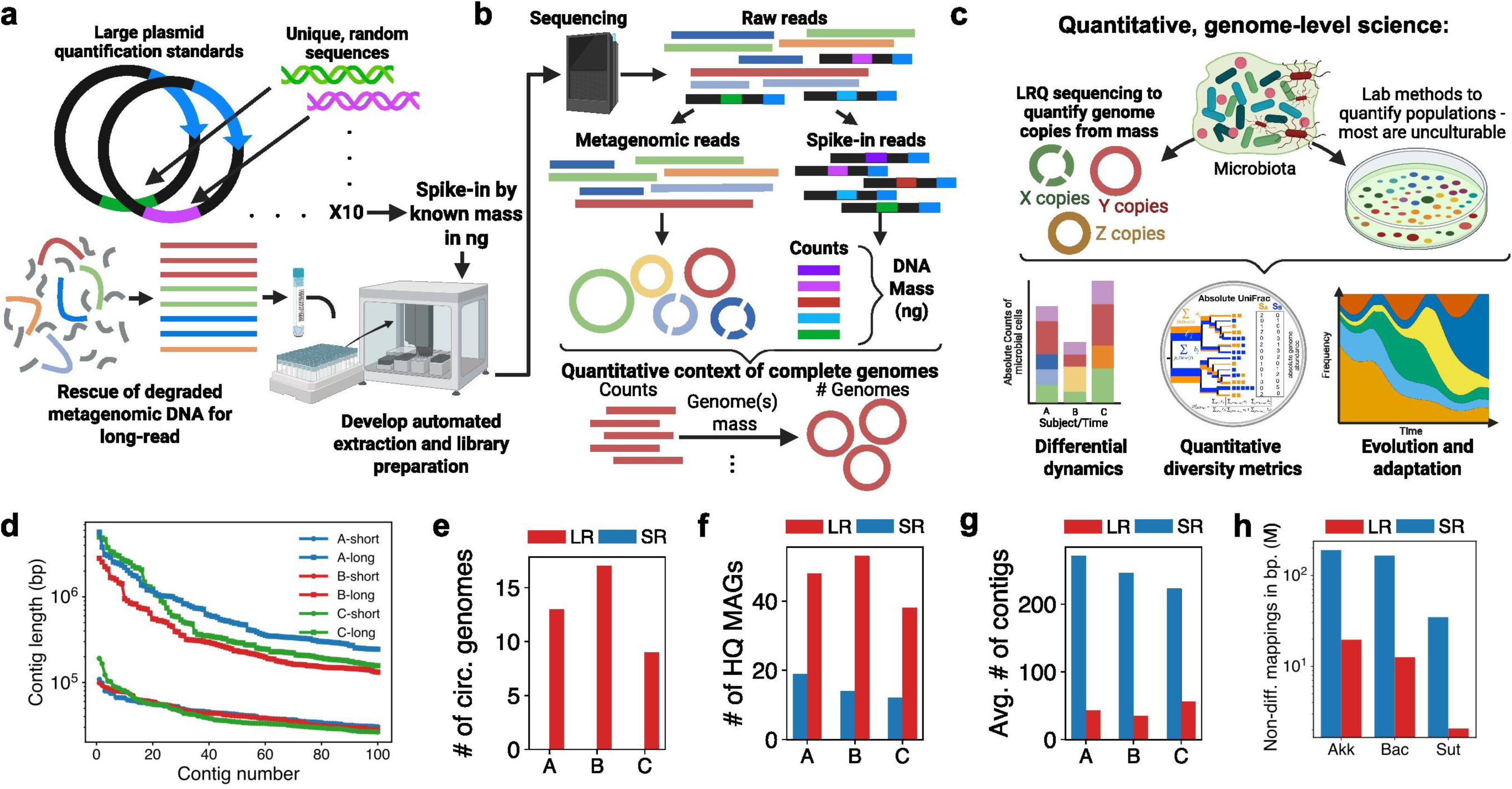
**a** Overview of the quantitative long-read metagenomics pipeline from DNA preparation. Large plasmids are constructed with unique, randomly generated insert sequences, and are subsequently sheared to the size range of long-read length. Commonly degraded metagenomic genomic DNA (gDNA) is rescued via size-selection and amplification. Quantification plasmids are pooled and spiked into sample gDNA and processed via a bespoke, automated library preparation workflow. **b** Prepared libraries are sequenced on the Revio platform, followed by bioinformatic processing of metagenomic and plasmid spike-in reads. The spike-in reads are used to generate an internal, per-sample relationship between DNA mass and read count. This relationship is then used to quantify long-read metagenome assembled genomes (LR-MAGs) which are assembled from the metagenomic reads. **c** Schematic detailing the advantages of quantitative genome-level science. Long-read sequencing allows the retrieval of circular or high-contiguous genomes, including unculturable microbes. Our quantification method provides absolute abundances of these LR-MAGs, enabling quantitative analysis of differential abundance, diversity, and evolution using absolute abundances instead of relative abundance. **d** Short-read and long-read comparisons of matched stool samples from volunteers showing the top 100 longest contigs from each approach. For short-read data, a non-linked read metagenomic assembler (metaSPAdes) was used on TELL-seq libraries. **e** The number of circular genomes and **f** HQ MAGs generated via assembly for each approach. **g** The average number of contigs per MAG from each approach. **h** The number of non-differentiable mappings for closely related reference genomes ∼(99.5% ANI) for each approach. Akk, Bac, and Sut correspond to *A. mucinophila*, *B. uniformis*, and *S. wadsworthensis A*, respectively. Panels A-C are created in BioRender. Din, O. (2025) https://BioRender.com/.

To investigate whether long reads provided the assembly contiguity and mapping specificity needed for strain-level inference, we compared assembly and mapping accuracy against shotgun metagenomics. We find that long-read metagenomics generates assemblies orders of magnitude longer than short-read assemblies (Fig. 1d). We also found dramatically more high-quality metagenome-assembled genomes (HQ-MAGs) with long-read sequencing (Fig. 1e-f), and obtained complete circular metagenome-assembled genomes (cMAGs) only with long-read. The average isolate genome size is ∼4.3Mb ^32^ thus, based on the achievable contig lengths, short-read MAGs are likely highly fragmented. Accordingly, our short-read MAGs on average consist of hundreds of contigs (Fig. 1g). Additionally, when comparing identical matches to closely related genomes, we observe that short-read data cannot differentiate highly similar genomes (e.g., 99.5% ANI) (Fig. 1h). These findings demonstrate specific advantages of our long-read metagenomics workflow over short-read approaches that enable assembly and discrimination at the strain-level, while providing a suitable springboard to develop quantification approaches.

To develop a long-read quantification approach that is portable and community-agnostic, we utilized a plasmid-based ‘synDNA’ framework which uses randomly generated inserts for read-mapping-based identification ^24^. Relative to the NCBI non-redundant nucleotide sequence database, and using blastn ^33^, all hits for these inserts have <50% query alignment ratio, and all hits with >15% query alignment ratio are <80% sequence identity (Extended Data Table. 1; Retrieved data May 03, 2026). To enable this approach for long-read sequencing, we cloned 10 unique synthetic fragments into ∼17.5-kb plasmids that shear within the High-Fidelity (HiFi) read-size range and spiked them into each sample at known masses (Fig. 2a). We found that plasmid DNA was more resistant to shearing than metagenomic DNA, so we measured the sheared fraction and used it to calculate the effective standard mass incorporated into each library (Extended Data Fig. 1a-d). After stool-DNA spike-in and sequencing, insert coverage was distinguishable from plasmid backbone coverage (Fig. 2b) and scaled linearly with plasmid input above 0.001 ng/uL (Extended Data Fig. 1e-f). We therefore used two plasmid-insert combinations per concentration across 0.001-10 ng/uL. Per-sample correlations between insert mass and read count, which also tracked coverage depth (Extended Data Fig. 1g), provided the calibration used to estimate target genome DNA mass (Fig. 2c-d). Reads derived from the plasmid backbones were removed before microbial analyses (see Methods). Thus, LRQ establishes a sample-specific relationship between known DNA mass and recovered long-read signal that is agnostic to community composition.

**Figure 2:**
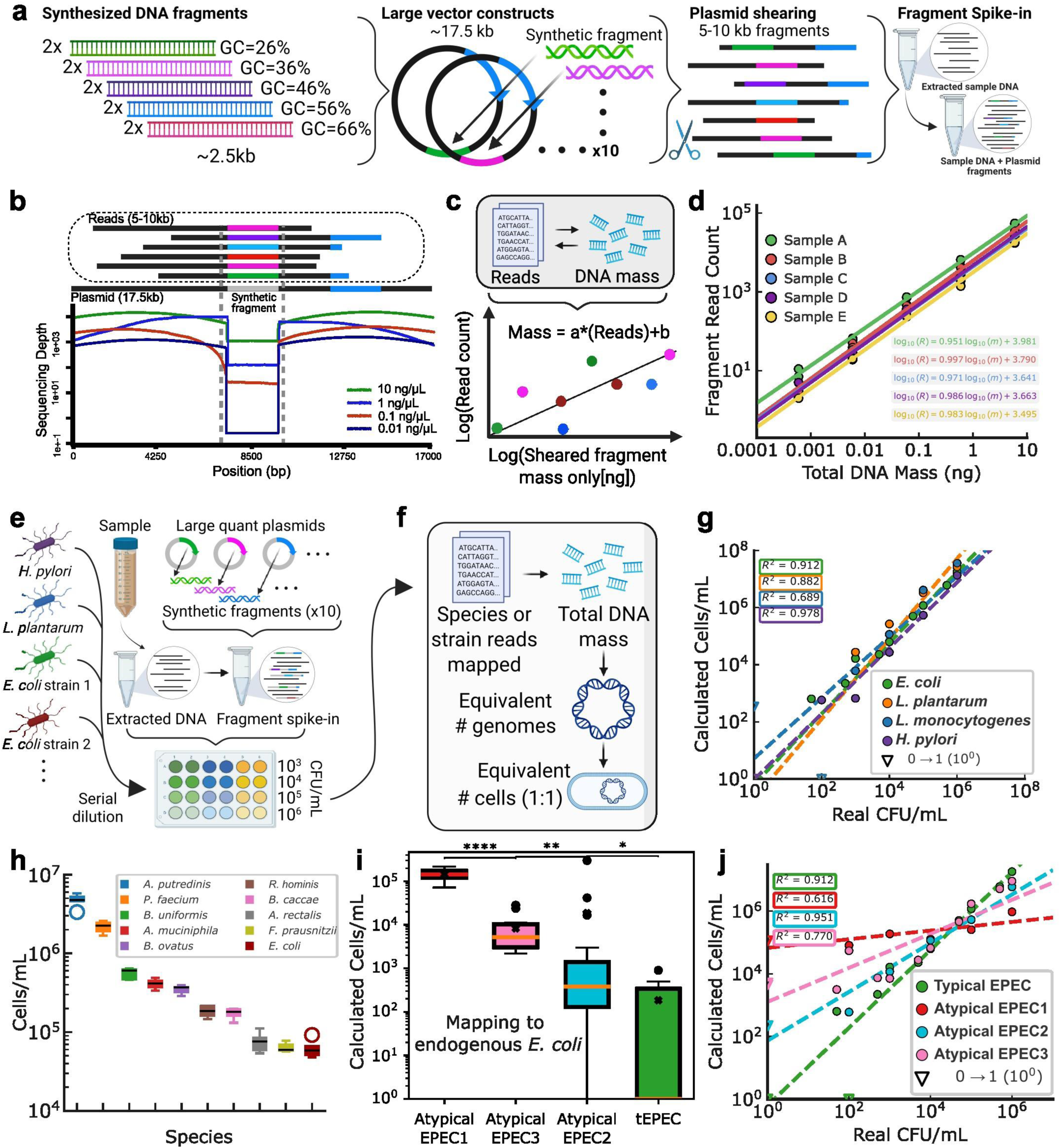
**a** Schematic outlining the design approach for constructing synthetic DNA fragments and incorporation into large plasmid backbones, followed by shearing and spike-in into gDNA samples. **b** Coverage plot showing the ability to discriminate different levels of spiked-in synthetic fragment DNA. **c** Schematic detailing the relationship used for relating DNA mass to sequenced reads. **d** Experimental validation of a linear dynamic range between fragment mass and read count, both log-transformed, at various spike-in levels into a stool sample. The results are consistent between various samples. R^2^ specified for each Sample/Subject on plot. **e** Experimental design of bacterial spike-in experiments to validate the ability of the method to accurately quantify bacterial absolute abundances in metagenomic samples. **f** Schematic detailing how the reads are used to generate a relationship with the amount of fragment DNA spike-in mass (see Methods). **g** Plot showing the validation of the quantification approach to accurately estimate bacterial absolute abundance compared to real CFU measurements. **h** Quantification of various microbial species in 5 technical replicates of the stool sample, demonstrating precision in measurements. **i** Plot showing the multi-mapping of endogenous *E. coli* against the genomes of the EPEC strains in one of the technical development stool samples (****P ≤ 1 × 10⁻⁴, **P ≤ 0.01, and *P ≤ 0.05; two-sided Mann–Whitney U tests with Holm correction for multiple comparisons, N=15-20 per group, 70 total across comparisons, U = 214-300 for these comparisons). **j** Spike-in experiments with the *E. coli* strains from **i** demonstrating that the long-read quantification method can discriminate closely related strains based on existing background. Panels A-C are created in BioRender. Din, O. (2025) https://BioRender.com/.

We next asked whether this internal calibration could recover microbial absolute abundances from stool. DNA from technical-development stool slurries was spiked with plasmid standards and bacterial strains diluted to 1 x 10^2-1 x 10^6 CFU/mL (Fig. 2e). Reads mapping to each strain genome were converted to genomic DNA mass using the per-sample calibration, then to genome equivalents by molecular weight and genome length (Fig. 2f; see Methods). Assuming a 1:1 ratio between genomes and cells, LRQ estimated cell numbers for four Gram-positive and Gram-negative GI-relevant species: *E. coli*, *L. plantarum*, *L. monocytogenes*, and *H. pylor*i (Fig. 2g). Estimates were slightly higher than culture inputs, consistent with rapidly dividing cells carrying more than one genome copy^34^. Quantification extended down to ∼100-1,000 cells/mL, corresponding to ∼1% coverage breadth (Extended Data Fig. 1h), and five stool technical replicates produced consistent species-level abundance estimates (Fig. 2h).

To determine the ability of LRQ to discriminate cell numbers at the intra-species level, we analyzed the quantitative sequencing approach on a set of microbial CFU dilutions with various enteropathogenic *E. coli* (EPEC), covering a wide array of pathogenic *E. coli* strains characterized by different phylogroups, antigens, and virulence factors. These included a typical EPEC (tEPEC) strain and three isolated atypical EPEC strains with varying levels of intra-species identity between them (Extended Data Fig. 1i). We observed significantly different levels of background burden characterized by reads mapped from endogenous *E. coli* (Fig. 2i), resulting in orders of magnitude differences in the calculated cell numbers for different strains. Upon microbial spike-in of these strains at different levels, the relationship between experimentally measured CFU counts and estimated cell numbers depended on the general background observed for each strain within the sample (Fig. 2j), showing that even within a complex sample, this method can achieve strain-level discrimination. Without any discrimination, we would expect background mapping and strain spike-in quantification results to be largely comparable between these different *E. coli*. Furthermore, we found that at 1 x 10^6^ CFU/mL spike-in of tEPEC, we could accurately recover the entire genome by metagenomic assembly, compared to a reference genome generated from a cultured isolate (Extended Data Fig. 1j). These observations show that long-read context can distinguish closely related genomes within a species, enabling strain-specific abundance measurements that would be masked in species-level analysis.

While recent work has been done to enable the use of low input DNA for long-read sequencing^35^, clinical and longitudinal microbiome studies often depend on stored specimens, creating a practical barrier for biology-forward long-read analysis because of DNA degradation^36,37^. Degradation is exacerbated by freeze-thaw cycles, commonly encountered when samples are transported or repeatedly used for experiments (Fig. 3a, left). Degraded DNA contains many small fragments that are preferentially incorporated into the library, resulting in significantly lower library mass and read length (Fig. 3a, top). To apply our quantification protocol to degraded metagenomic samples, we developed an amplification protocol that selectively removes short DNA fragments and subsequently amplifies long fragments (Fig. 3a, bottom). To verify that this approach did not introduce substantial biases, we extracted stool DNA from five individuals for technical development, and processed them both with and without the amplification protocol as described in Methods (Extended Data Fig. 2a-d).

**Figure 3:**
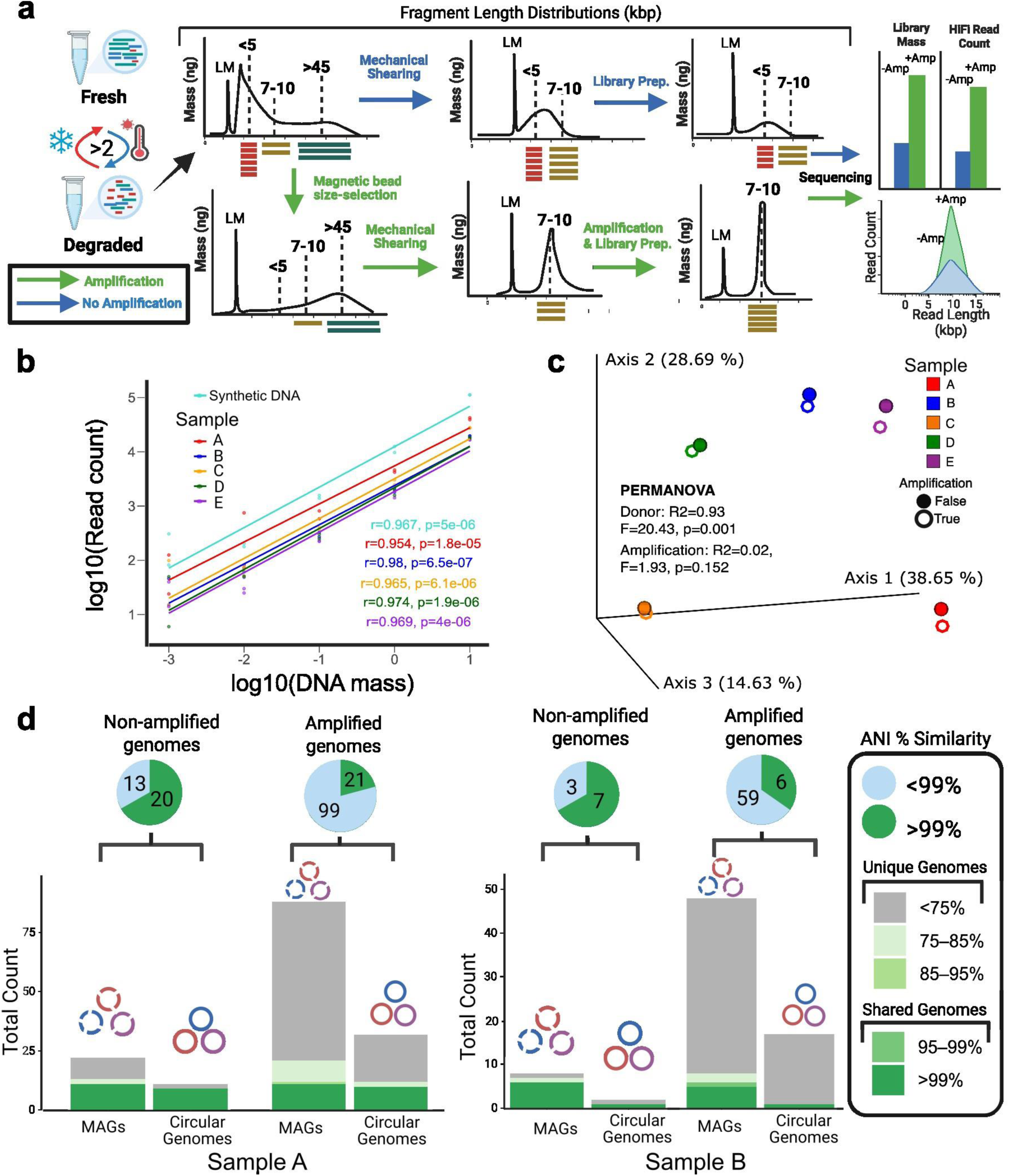
Rescuing low quality genomic DNA (gDNA) with size-selection and whole genome amplification. **a** Flowchart illustrates that combining size-selection with whole genome amplification greatly improves the library mass, HiFi read count and read length of degraded gDNA. **b** Relationship between input concentration and read output for ten plasmids carrying synthetic DNA sequences spanning a range of GC contents, each spiked into five human fecal samples and processed separately. **c** Principal Coordinates Analysis (PCoA) of Bray–Curtis dissimilarities for paired fecal samples from five human stool samples, comparing samples processed with and without size-selection and amplification R^2^ values specified on panel. **d** Number of high quality MAGs (completeness-4*contamination > 50) and circular genomes recovered under each method. Bars are colored according to the Average Nucleotide Identity (ANI) between genomes reconstructed with the two approaches. Panels A was partly created in BioRender. Din, O. (2025) https://BioRender.com/.

Size-selection with whole-genome amplification rescued degraded gDNA. In paired fecal samples subjected to repeated freeze-thaw cycles, amplified samples started from less input DNA (25 ng versus 600-900 ng) yet yielded higher library masses (22.7-84.8 ng versus 8.3-22.3 ng), more reads (0.92-4.31 million versus 0.08-0.60 million), and ∼3-10x higher total HiFi yield (Fig. 3a, right; Extended Data Table 2). Mean read lengths were comparable (Extended Data Fig. 2e-f). Ten synthetic plasmids spanning a range of GC contents retained strong input-output linearity across samples (r = 0.95-0.98, p < 10^−5^; Fig. 3b), and community composition remained primarily sample-driven rather than processing-driven by PERMANOVA (R^2^ = 0.02, p = 0.152; Fig. 3c). Amplification increased recovered MAGs (36-137 versus 18-54) and circular genomes (36-85 versus 6-40), with most paired genomes showing >99% ANI (Fig. 3d; Extended Data Table 2). Thus, size-selection and amplification enabled quantitative long-read recovery from degraded, low-input samples.

To evaluate LRQ in a biologically relevant setting, we applied it to a long time-series dataset of inflammatory bowel disease (IBD) from a colonic Crohn’s disease (CCD) patient, which is a biologically distinct from, and understudied relative to, ileal Crohn’s disease (ICD)^38^. This dataset captures longitudinal data before and after a sigmoid colectomy, the primary site of disease, which was associated with a shift from high inflammation to clinical quiescence (Fig. 4a, top)^39,40^. The dataset contains longitudinal measurements of inflammatory markers throughout the timeseries, providing information about various mechanisms that shape the microbial landscape of the gut (Fig. 4a, bottom)^41^. Because longitudinal datasets typically possess an inherent trade-off between temporal depth and microbial resolution, driven by short-read identification of high-level taxonomy or manual isolation of culturable microbes, application of LRQ to this time-series presented a deep proof-of-principle of whether LRQ could connect strain-level absolute abundance, host physiology and microbial evolution in the same samples^42–45^.

**Figure 4:**
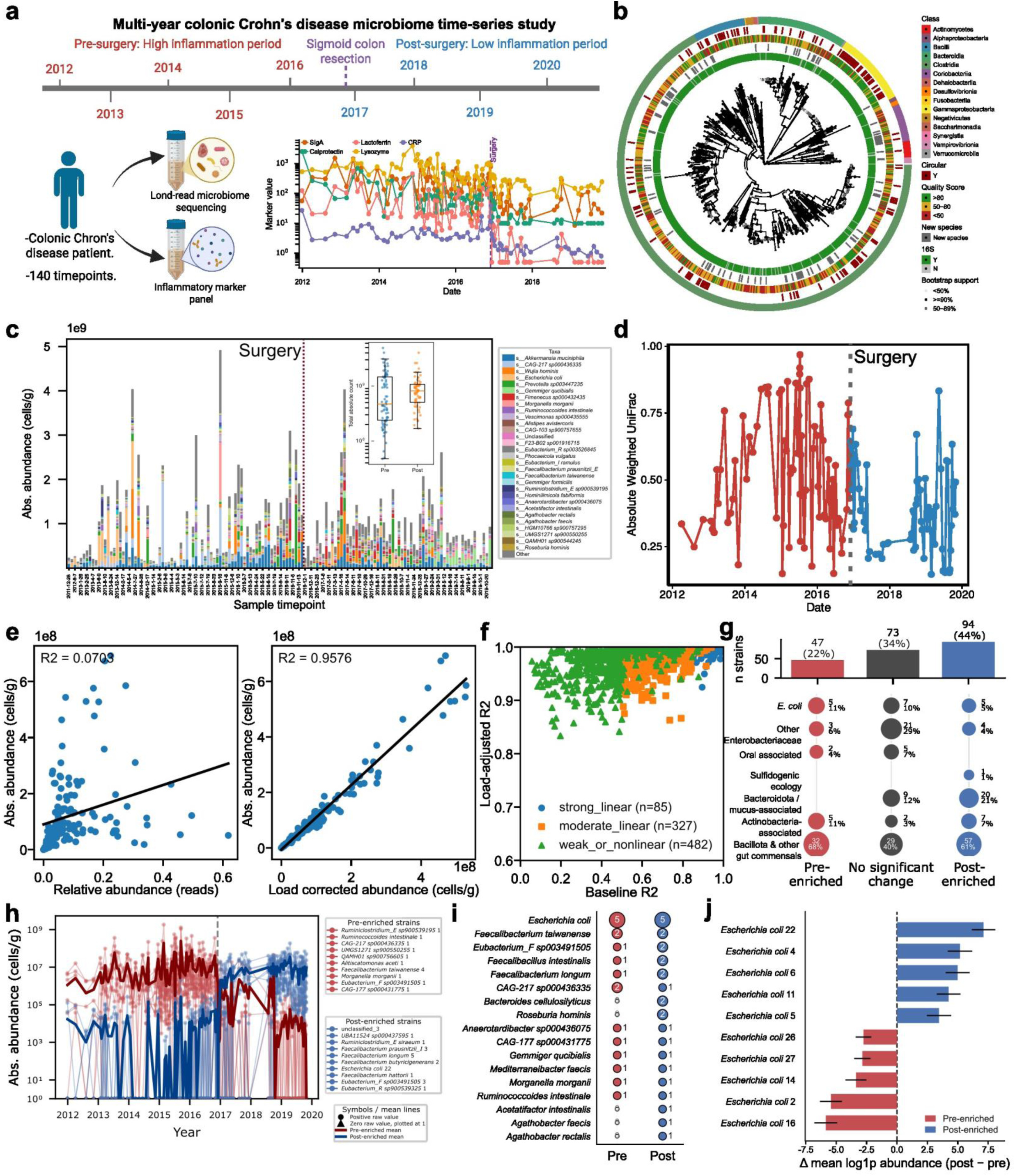
**a** Overview of a multi-year study of a colonic Crohn’s disease patient (CCD). The analyzed dataset spanned from late 2011 to late 2019, approximately 8 years. The patient experienced a period of high-inflammation during active disease, prior to a sigmoid colectomy in 11/2016, after which symptoms went into clinical remission (top). Whole stool samples were collected throughout the study for microbiome analysis, in addition to measurements of fecal inflammatory marker levels (bottom left). Marked recession of inflammation can be observed from the trajectories of five distinct inflammatory markers: calprotectin, lactoferrin, secretory IgA, lysozyme, and CRP (bottom right). We analyzed a total of 140 samples in this study. **b** Tree of all LR-MAGs generated from the 140 samples which were at least >50% completeness and <10% contamination. Various tracks show LR-MAGs by taxonomic class, circularity from assembly, overall quality score, presence of 16S, and whether it is identified as a new species. **c** Taxa barplot of all the samples in the time series scaled by the absolute quantification within each bar. Inset: plot of total absolute count from each sample in the pre and post surgery periods (*P < 0.05, Levene test on variance, W≈5.08, N=140 samples, with N=78 pre-surgery and N=62 post-surgery samples). **d** Absolute weighted UniFrac of time-adjacent samples (timepoint-to-next) across all samples in the timeseries. Red and blue indicates pre- and post-surgery periods, respectively (*P < 0.0005, Mann-Whitney U test for N=139 time-adjacent intervals, where the pre-surgery N=77 and the post-surgery N=62, with W statistic = 3322). **e** Relationship between calculated absolute abundance (cells/g) and reads-based relative abundance (mapped reads/total reads) for *A. mucinophila* across all samples (left), and after load adjustment (read-based relative abundance x total per-sample absolute abundance (right). **f** Relationship between the R^2^ value for the baseline relative abundance and load-adjusted correlations with absolute abundances across all LR-MAGs. **g** Differential abundance of all LR-MAGs >90% completeness, showing pre- and post-enriched LR-MAGs across the 140 sample timepoints, as well as those not significantly enriched in either period (top). General breakdown of the type of microbes in the set of LR-MAGs (bottom). **h** Absolute abundance trajectories of the top pre- and post-surgery enriched LR-MAGs (red and blue, respectively). Solid lines represent mean trajectories for either group. **i** Differential abundance of LR-MAGs across different species pre and post surgery. **j** Differential mean log1p abundance of *E. coli* LR-MAGs which are either pre or post surgery enriched (red and blue respectively). Error bars denote the combined standard error from the pre- and post-surgery windows (N=78 and 62 samples for the pre- and post-surgery windows, respectively). Significance for differential abundance was determined with an empirical block-shuffle permutation test; using BH-FDR q < 0.05. Panel A is created in BioRender. Din, O. (2025) https://BioRender.com/.

LRQ sequencing of 140 time-series samples yielded 5,453 MAGs, including circular contigs, at >50% completeness and <10% contamination (Fig. 4b; Extended Data Fig. 3a). These included 1,140 circular complete genomes and 168 single-contig MAGs. After de-replication at 99.5% ANI to achieve strain-level MAGs and selection of the highest-quality representative from each cluster^31^, we obtained 934 LR-MAGs, including 138 complete circular MAGs and 29 additional single-contig MAGs; representative genomes had an average quality score ≥50, consistent with literature thresholds^46^. Absolute quantification revealed that total microbial load varied by two orders of magnitude and became significantly less variable after surgery (Fig. 4c; Extended Data Fig. 3b). Absolute weighted UniFrac^47,48^ of time-adjacent samples showed a higher number of large phylogenetic shifts before surgery than after surgery (Fig. 4d), indicating higher strain-level turnover during high inflammation.

To explore differences between absolute and relative abundance, we analyzed *A. muciniphila*, one of the most persistent microbes in the time-series, which was represented by a single LR-MAG. We found that the relationship was not linear across the samples (Fig. 4e, left); however, a linear relationship could be achieved with a load-corrected abundance (total per-sample absolute abundance x relative abundance) (Fig. 4e, right). We then analyzed these correlations across all LR-MAGs and observed that most were weak or non-linear in their relationship, where load correction was needed to linearly relate the read-based relative abundance with absolute abundance (Fig. 4f). These results indicate that relative abundance values are not always linearly proportional to absolute abundance data.

Although some efforts have been made to study the differential abundances of microbial taxa before and after surgery for IBD, study of microbial strains in this context is still challenging due to technical limitations of relative abundance data and highly fragmented genome assemblies from short-read approaches^40,49^. Among 214 de-replicated LR-MAGs with >90% completeness, nearly half were post-surgery enriched, 22% were pre-surgery enriched, and almost one-third did not significantly change, suggesting distinct ecological responses to host state (Fig. 4g, top). Within *E. coli*, which has been strongly implicated in Crohn’s disease^50,51^, we find that LR-MAGs are distributed almost evenly across these three enrichment states, while other Enterobacteriaceae generally did not change pre- or post-surgery (Fig. 4g, bottom). Two potentially oral-derived microbes are enriched pre-surgery, *P. micra* and *P. stomatis*, which are associated with various oral diseases and potentially cancer^52–54^. Other broad groups of microbes were found to be mostly enriched post-surgery (Fig. 4g, bottom). Absolute abundance trajectories differed by one or more orders of magnitude between pre- and post-surgery enriched LR-MAGs, which would be unobservable from relative abundance alone (Fig. 4h). We also found that differential intra-species enrichment extended beyond *E. coli*, including another IBD-implicated pathogen, *M. morganii*, and two species of *Faecalibacterium*, which are generally considered anti-inflammatory in the broad IBD context (Fig. 4i and Extended Data Fig. 3c-d)^55–57^. Differential abundance of *E. coli* LR-MAGs further demonstrated the significant intra-species count differences that can be observed (Fig. 4j). Furthermore, intra-species turnover and persistence can vary dramatically between high and low inflammatory regimes, further supporting the view that these LR-MAGs are distinctly affected by their environment (Extended Data Fig. 3e-f). These results show that intra-species resolution is necessary to identify associations that would have otherwise been missed at higher taxonomic levels, and that a single species should not necessarily be viewed as strictly ‘disease-associated’ or ‘health-associated’.

Because our study includes a unique combination of time-series data on *de novo* assembled LR-MAGs and dense sampling of host inflammation, LRQ presents a unique opportunity to compare the dynamics of microbial genomes against specific inflammatory markers over multiple years. For this dynamical analysis, we endeavored to use dynamical comparisons uniquely enabled by absolute abundance data^58^. To determine which LR-MAGs were tracking unique inflammatory markers, we analyzed correlations during active inflammation (pre-surgery period) across the following dynamical properties: co-movement, lagged, detrended, and change tracking (Fig. 5a and Extended Data Fig. 4a-b). We found many microbes that positively or negatively tracked the different markers, with a few mixing positive and negative correlations. Most of the negative trackers correlated with lysozyme^59^, which is consistent with its mechanism of action that directly targets bacterial cell walls, while most positive trackers were associated with either calprotectin or lactoferrin (Fig. 5b-c). When considering intra-species differences, we observed that various species can have multi-marker trackers, in some cases with both positive or negative correlation (Fig. 5d). *E. coli* possessed the most differential trackers across all inflammatory markers, with almost equal numbers of positive and negative marker trackers (Fig. 5e-f). Therefore, LRQ enabled the finding that LR-MAGs can dynamically track specific host inflammatory markers, and that intra-species LR-MAGs can exhibit opposing tracking behavior.

**Figure 5:**
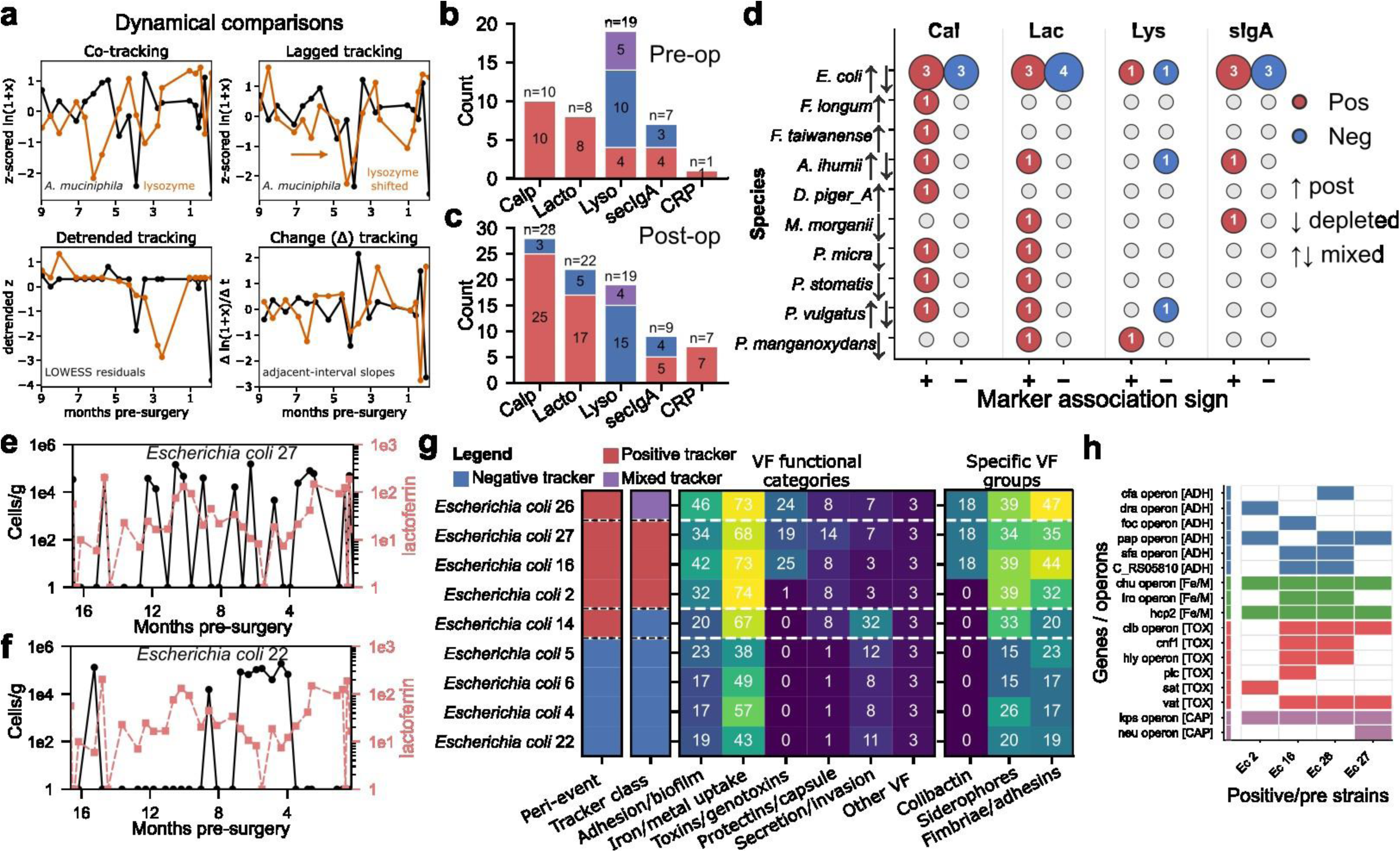
**a** Summary of dynamical comparisons performed between microbial genomes and inflammatory marker trajectories. Tracker significance was assessed using an empirical block-shuffle null model; marker–feature pairs were called significant at P_emp_ < 0.05 across the 140 timepoint samples (see Methods). **b** Breakdown of positive and negative marker trackers (red and blue, respectively) for pre, and **c** post, surgery. Purple indicates LR-MAGs that have mixed positive and negative correlations within the dynamical comparisons. **d** Breakdown of positive and negative marker trackers across various microbial species. LR-MAGs can possess positive or negative correlations across different markers. Cal, Lac, Lys, and sIgA refer to calprotectin, lactoferrin, lysozyme, and secretory IgA, respectively. **e-f** Trajectories of cell count and lactoferrin for one of the **e** positive and **f** negative marker trackers. **g** Heatmap showing the presence of virulence factors (VFs) at the level of functional categories (center) and specific groups (right) for both the positive and negative *E. coli* inflammatory marker trackers. **h** VF genes and operons only present in the pre-enriched, positive tracker *E. coli* compared to the post-enriched, negative trackers.

Calprotectin, lactoferrin, lysozyme, and secretory IgA have negative effects on microbial growth by modulating metal availability, lysis, and adherence, respectively^60,61^. We therefore hypothesized that positive inflammatory marker trackers, particularly those from *E. coli,* were potentially pathogenic LR-MAGs which were able to tolerate or benefit from the inflammatory environment^62^. We observed that most positive *E. coli* trackers were also pre-surgery enriched, while most negative trackers were post-surgery enriched (Fig. 5g, left). Analysis of the absolute abundance trajectories of *E. coli* LR-MAGs showed that variable strain-level blooms materialize across the time series broadly grouped between pre-surgery enriched/positive trackers and post-enriched/negative trackers (Extended Data Fig. 4c-f). Furthermore, episodic strain dominance is not always consistently associated with inflammatory-marker tracking or pre- and post-surgery enrichment when observing non-trackers and non peri-event changers, whose presence is more homogeneously spread across the time-series (Extended Data Fig. 4g-i). To determine whether positive trackers had more pathogenic properties than negative trackers, we mapped these LR-MAGs to the Virulence Factor Database (VFDB) and assessed the broad functional categories of virulence factors (VFs) present in either group. We found that the positive tracker possessed higher numbers of biofilm, metal uptake, toxin, and capsule VFs compared to the negative tracker group (Fig. 5g, center). We further found that within these categories, the most enriched groups of VFs were genotoxins, siderophores, and fimbriae and adhesin genes (Fig. 5g, right, and Extended Data Fig. 5a). Delineating these VF groups to specific genes and operons present in the positive tracker group, and not the negative trackers, we found that prominent and emerging virulence factors, such as the colibactin-producing *clb* operon, the a-hemolysin producing *hly* operon, the immune-evasion *kps* operon, and the host-iron utilizing *chu* operon (Fig. 5h)^63–66^. Some of these VF’s are sparsely present in other groups of *E. coli*, however, such as non-tracker/peri-event changers and other pre- or post-enriched *E. coli* (Extended Data Fig. 5b). These results demonstrate that certain LR-MAGs that positively track with calprotectin and lactoferrin have more VF’s that can not only counteract these markers by likely increased metal uptake, but also have a unique combination of other VF’s that are known to be broadly pathogenic in various disease contexts. Thus, absolute abundance data generated by LRQ enabled the determination of *E. coli* LR-MAGs with high pathogenic potential based on dynamical correlations of absolute abundance dynamics and inflammatory markers.

Next, we considered adherent-invasive *E. coli* (AIEC) strains previously isolated from Crohn’s disease patients. Specifically, we examined LF82 and NRG857c^67^, observing high ANI with the *E. coli* pre-surgery enriched/positive-tracker group (Extended Data Fig. 5c). We also found near-complete identity between *E. coli* 27, a single-contig assembly, and a previously sequenced AIEC-like isolate from this same patient^39^, where the LR-MAG was of significantly higher quality than the short-read assembled isolate genome (Extended Data Fig. 5d-e). Additionally, we also analyzed an LR-MAG from another species, *F. prausnitzii*, and found that much of the genome possessed high ANI compared to a public database reference genome (Extended Data Fig. 5f). These results showed that the *E. coli* pre-surgery enriched/positive-tracker group assembled from LRQ were very similar to known AIEC isolates from Crohn’s disease patients.

Because LRQ recovers complete genomes and high-quality MAGs, in addition to their absolute abundances over time, they can be used to enable the analysis of evolutionary dynamics over time. To explore these dynamics, we developed a hybrid graph-probabilistic haplotype reconstruction pipeline called Strainphase, which links localized genetic variations into contiguous, long-range haplotype tracks (Fig. 6a, left). Tracks with shared consensus across timepoints were clustered into stable lineages, serving as our primary unit of evolutionary observation (Fig. 6a, center and top right). The mean size of each phase block was 22kb, showing the ability of long-reads to capture haplotype information across many genes (Fig 6a, bottom right). To determine what functions were under selection, we mapped SNV-bearing genes within sweeping lineages to curated KEGG pathway categories ^68–70^. Functional pathways targeted by sweeps were distinct across the peri-surgical transition, with pre-surgery high-inflammation samples enriched for pathways consistent with inflammatory stress adaptation and microbial turnover, including DNA replication, mismatch repair, homologous recombination, oxidative phosphorylation, and PTS-mediated nutrient uptake (Fig. 6b) ^71^. In contrast, post-surgery remission was enriched for sweeps in pathways more consistent with metabolic rebuilding and anaerobic gut function, particularly propanoate (SCFA) metabolism, amino acid and vitamin biosynthesis, suggesting host-state-dependent selection on distinct microbial ecotypes across the inflammatory transition (Fig. 6b)^72^. Analysis of the selective sweeps against the pre- and post-surgery periods across all of the LR-MAGs revealed a stark inflection at the point of surgery (Fig. 6c). Sweep midpoints heavily accumulate in the weeks immediately flanking the operation, consistent with the understanding that the surgery served as a selective bottleneck. We then visualized how the surgical intervention impacted strain diversity (lineages per LR-MAG) and lifespan, and found that lineage persistence and dominant-lineage turnover, as a function of the number of lineages per LR-MAG, differentially shifted across the peri-surgical transition (Extended Data Fig. 6a-b). *A. muciniphila*, which generally appeared to be a stable outlier compared to other species with respect to its absolute abundance, showed it carried a high number of lineages per LR-MAG and one of the longest mean lineage persistences. This indicated that its long-term presence may be maintained by a reservoir of persistent haplotypes, as opposed to inflammation-related blooms of differentially enriched strains. Taken together, these results demonstrate that LRQ can enable discovery of sweep events in gene-level windows functionally relevant to CCD and characterization of strain-level ecology throughout phenotypic windows.

**Figure 6:**
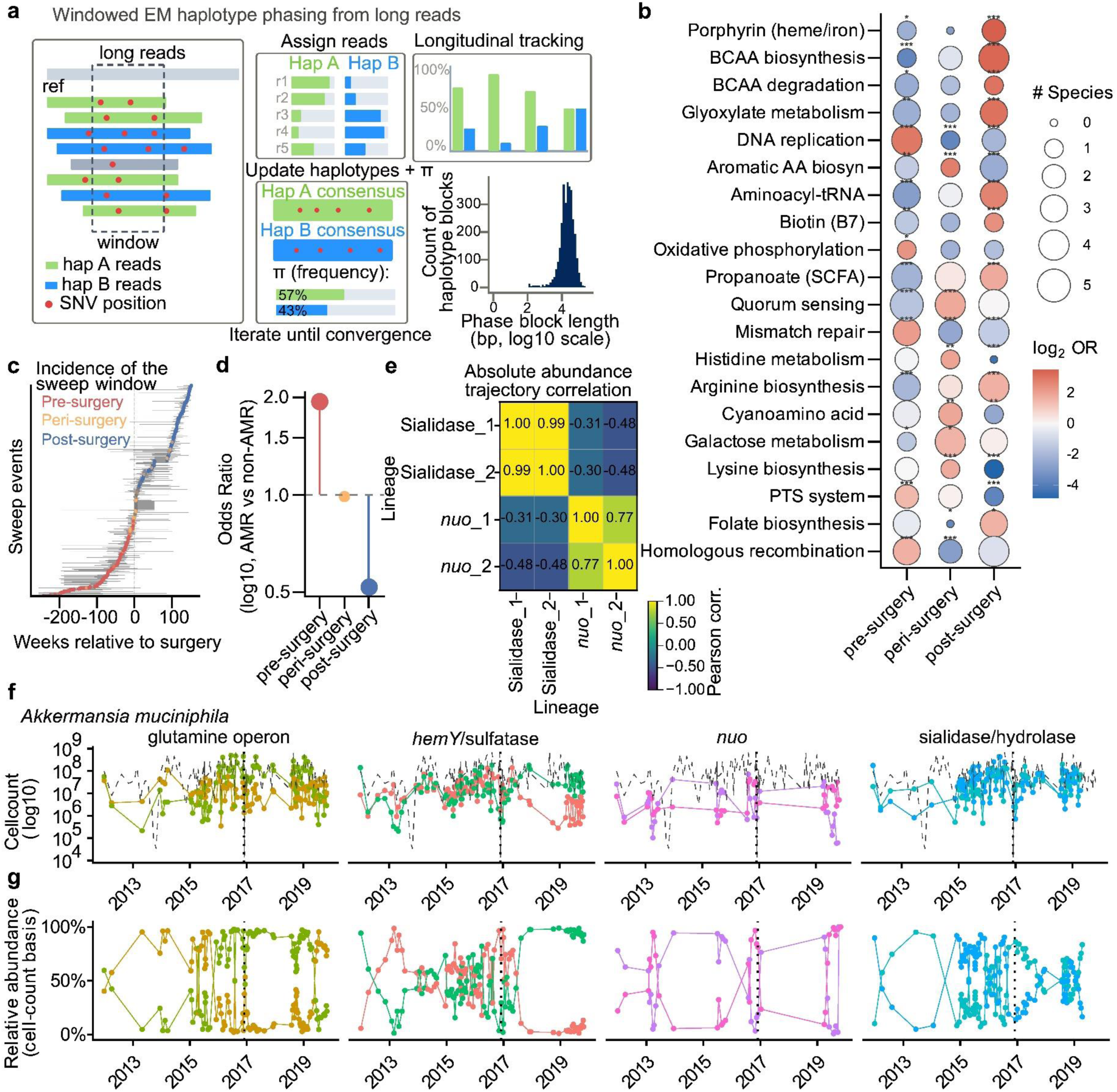
**a** Strainphase algorithm schematic. Aligned PacBio HiFi reads and variant calls are processed in overlapping windows to build read-overlap graphs, partitioned via community detection. Redundant haplotypes are merged, linked into contig-spanning tracks, and “rescued” across timepoints by matching low-weight reads to anchor haplotypes. These tracks are clustered into cross-sample lineages. **b** KEGG pathway enrichment by sweep timing. Categories with ≥10 gene events per window were tested by two-sided Fisher’s exact test (one-vs-rest within each timing window) on gene-event counts (pre-surgery n = 2,968; peri-surgery n = 1,014; post-surgery n = 1,419 KEGG-annotated gene events from sweeping lineages). Bubble fill, log₂(odds ratio) (blue = depleted, red = enriched); bubble size, number of distinct species contributing gene events to that cell (range 0–5). Asterisks, Benjamini–Hochberg–adjusted significance: *p_adj < 0.05, **p_adj < 0.01, ***p_adj < 0.001. **c** Sweep midpoint barbell plot. Each row is a detected sweep event (a haplotype-lineage trajectory transitioning between ≤20% and ≥80% relative abundance within a temporal window). Grey bars span the shortest transition window; colored points mark the 50%-abundance midpoint. Point color denotes timing: pre-surgery (red, n = 917 sweep events), peri-surgery (orange, ±4 weeks, n = 363), post-surgery (blue, n = 1,235); an additional 306 events with indeterminate timing are not shown. **d** AMR enrichment by sweep timing. Lollipop chart of odds ratios (log₁₀ y-axis, 95% exact CI) from two-sided Fisher’s exact tests on sweep-event counts (denominators as in b) for events whose annotated products using Bakta are involved in AMR. Pre-surgical sweeps are significantly enriched for AMR genes (OR = 1.94, 95% CI 1.46–2.57; BH p_adj = 2.7 × 10⁻⁵), post-surgical sweeps are significantly depleted (OR = 0.52, 95% CI 0.39–0.69; BH p_adj = 3.2 × 10⁻⁵), and peri-surgical sweeps fall on the null (OR = 0.99, p = 1.0). Dashed line, OR = 1. **e** Absolute abundance trajectory correlation among four *A. mucinophila* lineages. Pearson correlation coefficients (r) were calculated from log1p-transformed absolute abundance trajectories across n = 140 longitudinal samples. Two-sided Pearson correlation tests were used, with Benjamini–Hochberg FDR correction across all lineage-pair tests in the bin; displayed pairs are FDR-significant (q < 0.05). Effect sizes are Pearson’s r. **f-g** Dynamics tracked across four functionally annotated loci showing **f** absolute haplotype cell absolute abundances (log₁₀ CFU) over time and **g** relative haplotype abundance.

Given the subject’s early antimicrobial exposure (01/2012 - 02/2012) in the early period of the timeseries, we also tested whether AMR enzymes changed over time. We screened for genes associated with resistance to several drug classes and found that AMR enzymes were significantly enriched among pre-surgical sweeps, whereas peri- and post-surgical sweeps did not enrich AMR enzymes (Fig. 6d). AMR sweeps generally targeted beta-lactamase, efflux, and fluoroquinolone resistance genes (Extended Data Fig. 6c). Fluoroquinolone-class sweeps were especially consistent with the recorded ciprofloxacin exposure, while the broader AMR enrichment suggests that LRQ can connect clinical perturbations to strain-level evolutionary responses.

To illustrate lineage dynamics over time, we focused on *A. muciniphila*, a mucus-specialized gut symbiont whose mucin-degrading lifestyle, genomic diversity, and context-dependent effects on host physiology make it a particularly informative target for genome-resolved evolutionary analysis, especially because it maintained high total abundance across the entire time series, both pre- and post-surgery^73^. We used the fact that our quantification model relates DNA mass to sequenced read count to quantify the absolute abundances per haplotype, then extended this number as an approximation of the cell-abundances possessing that haplotype using a 1:1 assumption between a haplotype and a genome, or cell. We tracked haplotype dynamics across respiratory/redox homeostasis (nuo complex), host-glycan/sialic-acid foraging (sialidase), tetrapyrrole/heme biosynthesis (hemY), and nitrogen assimilation (glutamine operon) functional groups^74–78^. Some lineage groups track each other strongly within gene-groups, such as sialidase and nuo, but other lineages negatively track between gene groups (Fig. 6e). These two groups of lineages exhibited a reciprocal relationship, where generally only one group was present at any given time. Furthermore, when comparing the trajectories of the absolute abundances and relative abundances of the lineages, we found that the lineage abundances broadly correlated with each other even when sweep events were observed in relative abundance (Fig. 6f-g). We also observed similar lineage dynamics in other LR-MAGs (Extended Data Fig 7). These results indicated that some paired haplotypes are likely embarking on very similar ecological strategies as evidenced by highly correlated dynamics. Thus, we showed that LRQ-enabled quantitative tracking of LR-MAG haplotypes across time can reveal haplotype dynamics that would otherwise not be observable with relative abundance data or with short-read metagenomics.

## DISCUSSION

This study establishes LRQ as a framework for observing microbiome biology as absolute, strain-level population dynamics. For the last decade, short-read metagenomics has greatly expanded the genomic diversity in reference databases of microbes beyond specific loci, and almost a million microbial genomes have now been deposited in public databases ^79^. However, short-read assembled MAGs are often highly fragmented and prone to chimerism due to the short-sequence context ^80^. Fragmentation, sequence similarity between taxa, and ambiguous read assignment all undermine short-read approaches when combined with internal-standard quantification. LRQ overcomes these challenges by pairing long sequence context with internal standards that match the long-read length, while yielding many *in situ* microbial circular and/or complete MAGs.

The technical advance of LRQ is the integration of three components that have usually been separate: long-read genome recovery, internal absolute calibration and degraded-sample rescue. The plasmid standards provide a per-sample relationship between known DNA mass and recovered long-read signal, allowing abundances to be reported as genome-equivalent estimates rather than as relative profiles alone. Validation with Gram-positive and Gram-negative bacterial spike-ins, closely related E. coli strains, and stool technical replicates showed that this calibration can support strain-level quantification. The size-selection and amplification workflow further extends LRQ to degraded or low-input specimens while preserving spike-in linearity, opening the possibility of applying quantitative long-read metagenomics to historically banked clinical and environmental cohorts.

Compared with 16S/qPCR, flow-cytometric load correction, cellular spike-ins, short-read synthetic standards and long-read assembly alone, LRQ is distinguished by assigning internally calibrated abundance to individual strain-level genomes and haplotypes. This distinction is significant because the method represents not only a sample-preparation or assembly workflow, but a measurement platform that links genome identity, abundance and evolution in the same assay.

Applying LRQ to a multi-year time series from a CCD patient who underwent sigmoid colectomy, we generated over 5,000 LR-MAGs, approximately one-fifth of them circular complete assemblies, and we then de-replicated them at 99.5% ANI, which yielded strain-level representative genomes. Quantification across the time series revealed marked changes in microbial load variability with respect to surgery. Absolute-weighted UniFrac uncovered dramatic community shifts between the high-inflammation period and post-surgery remission. Differential abundance analysis on the highest-quality LR-MAGs revealed substantial intra-species variability across the peri-surgical transition, showing that LR-MAGs from the same species can follow markedly different trajectories with disease state.

*E. coli*, pathogenic strains of which are associated with IBD, yielded the highest number of LR-MAGs which were differentially enriched, where persistence and turnover were significantly altered depending on the enrichment group. Absolute abundances allowed us to discover dynamical relationships between microbial genomes and inflammatory marker levels, where a set of *E. coli* LR-MAGs were found to bifurcate into positive and negative trackers, respectively. Pre-surgery enriched, positive trackers were found to possess a significantly higher burden of VF genes that may potentiate immune evasion and encode genotoxins. This approach enables negative and positive tracking as methods to identify microbes that are susceptible to the effects of host anti-microbial strategies, and those which tolerate, or even benefit from, host inflammation.

LRQ also made it possible to generate long-context haplotypes and to detect their sweeps across the LR-MAGs. Sweep analysis identified pathogenic pathways and AMR genes which underwent selection in the pre-surgery, high-inflammation period. Our quantification model allowed us to determine the absolute abundances of each haplotype lineage across the time-series, showing that some lineages can correlate in absolute abundance while showing sweep-like behavior in relative abundance. We anticipate that the ability to quantify haplotypes within the microbiome will enable new types of evolutionary research that are impossible with relative abundance only.

This approach will be limited by the efficacy of existing extraction protocols, especially for organisms that are particularly difficult for DNA extraction with microbiome kits, such as fungi or microbial spores. Absolute abundance estimates will also be limited by effective genome copy-number per cell, as this may vary depending on the strain and growth state. Since the approach is sequencing-based, there will also be limitations based on detection of very low abundant microbes. Furthermore, presently achievable sequencing error-rates may also be a limiting factor for higher strain resolution via mapping approaches, where inter-strain ANI, depending on the definition of a ‘strain’^10^, could plausibly be upwards of 99.9%. Accurately discriminating these strains in the microbiome will require even longer and more accurate reads than those currently achievable with existing long-read sequencing technologies. Even with these constraints, Future work will pursue experimental and computational improvements to span the full range of inter-strain similarity. We anticipate that LRQ will extend microbiome science to strain-level resolution across diverse environments and enable comparable analyses across fields.

## METHODS

Methods attached as a separate document.

---

**Extended Data Fig. 1:**
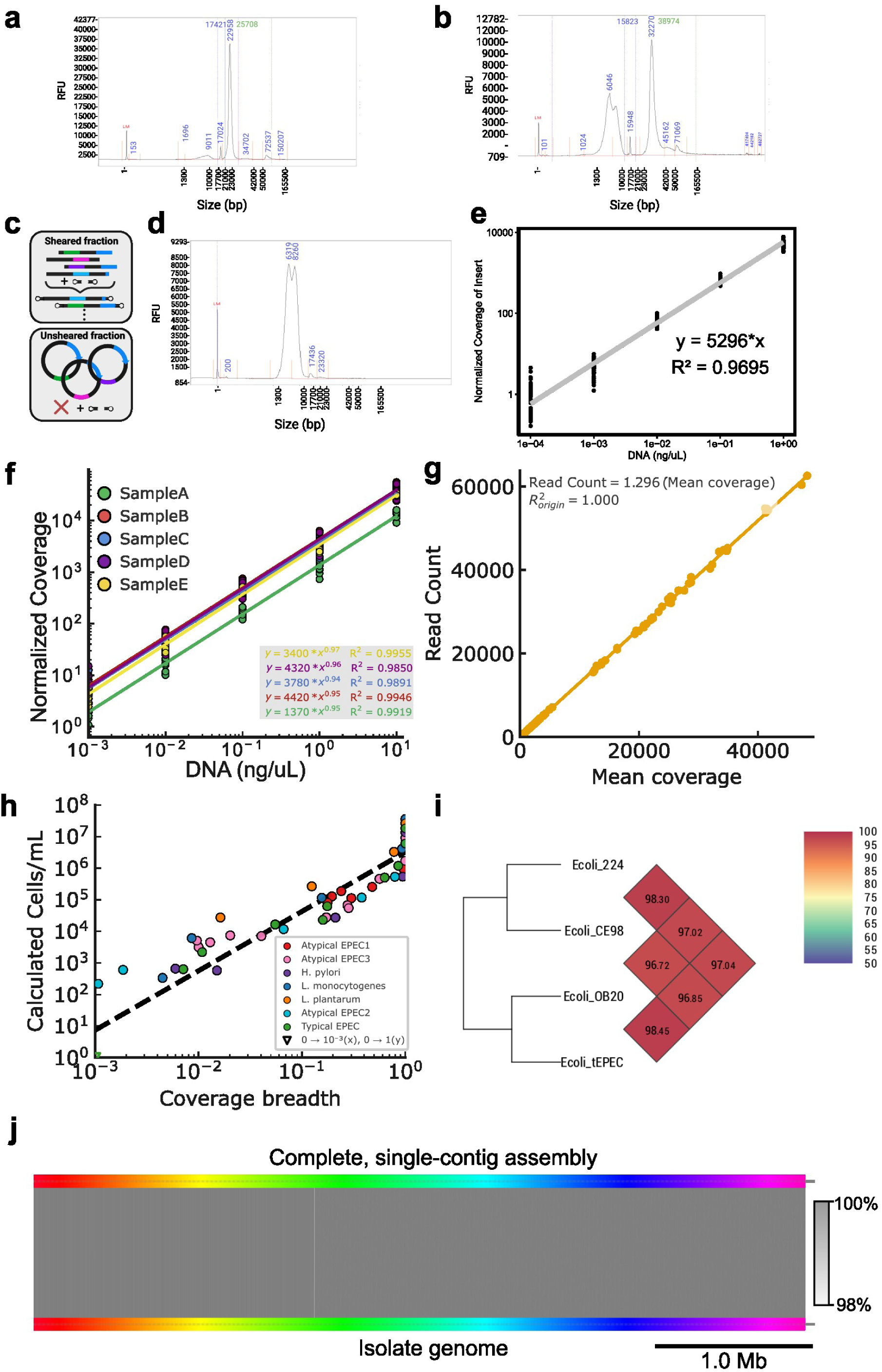
**a** Femto Pulse (Agilent) electropherograms of the 10 synthetic DNA-carrying plasmid pool before shearing and **b** after shearing. **c** Schematic of the sheared and unsheared DNA fractions. Sheared DNA allows adapter ligation and sequencing of fragments, whereas the unsheared fraction preserves circular molecules that may not be adapter-ligated and properly incorporated in the resulting library preparation. **d** Resulting PacBio HiFi library on the sheared plasmid pool. **e** Normalized coverage (to total bp sequenced per-sample) of defined synthetic DNA fragment sequences increases linearly with input DNA concentration, demonstrating quantitative recovery of the calibration molecule across the dilution series in volunteer stool samples. However, measurements become more variable at .0001 ng/uL. **f** Serial dilution of input DNA from five volunteer stool samples shows that normalized plasmid sequence (to total bp sequenced per-sample) coverage scales approximately linearly with input plasmid DNA concentration across several orders of magnitude. **g** Read count is tightly proportional to mean plasmid sequence coverage depth, indicating that mapped-read quantification is consistent with the underlying sequence coverage. **h** Estimated cell concentrations as a function of coverage breadth track the expected dilution series across multiple bacterial taxa. **i** Pairwise genome similarity via ANI among four closely related *E. coli* strains used for benchmarking strain-level discrimination; despite high nucleotide identity, the strains remain distinguishable at the genome level. **j** Whole-genome alignment between a complete, single-contig long-read assembly and the corresponding isolate genome for tEPEC shows near-complete colinearity and high nucleotide identity, demonstrating that the assembly approach can recover closed bacterial genomes. Gray colored lines between the two assemblies denote the level of identity (scale bar on right).

**Extended Data Fig. 2:**
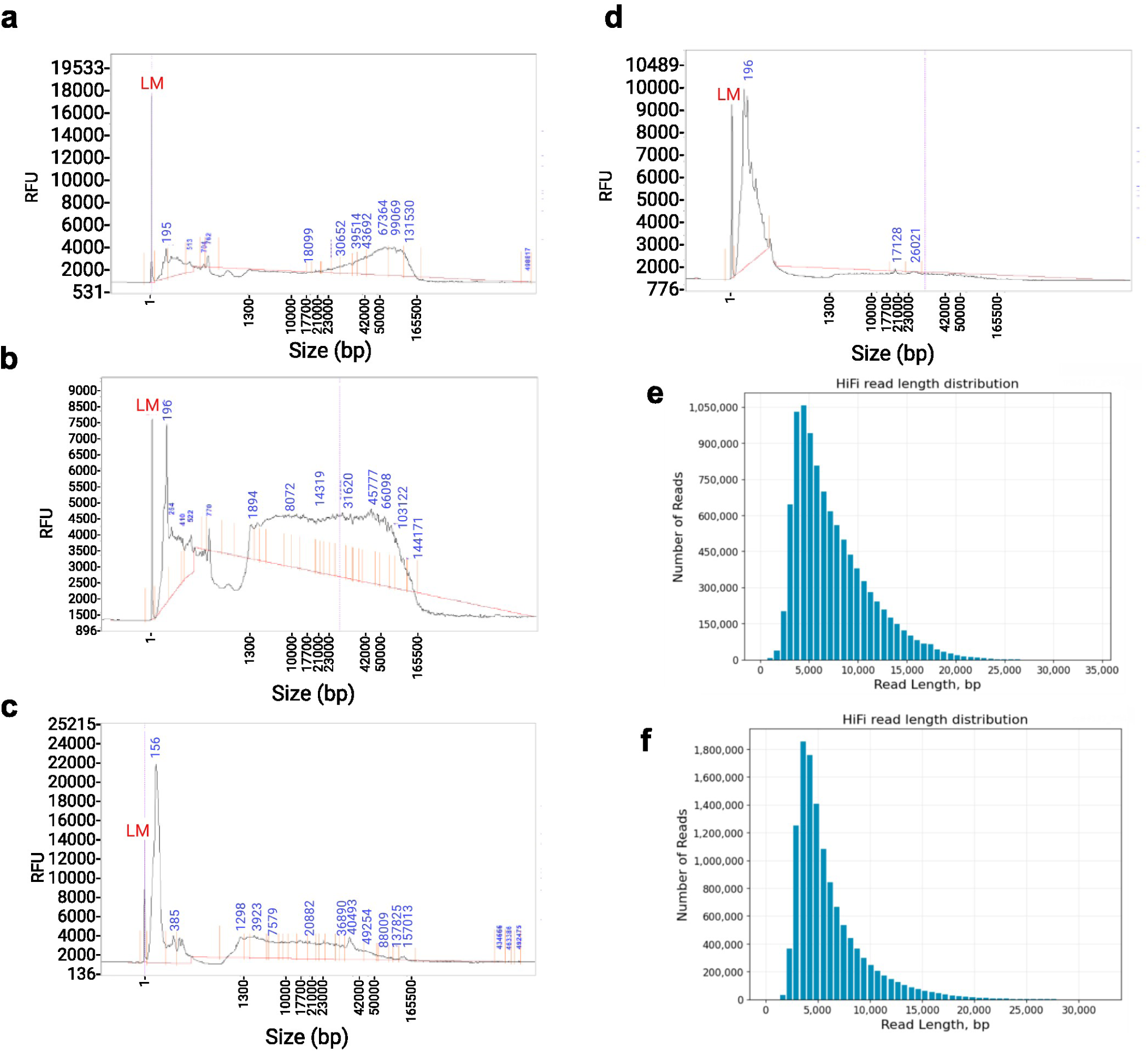
Femto Pulse (Agilent) electropherograms of genomic DNA extracted from human fecal samples. **a** Volunteer stool sample processed without multiple freeze–thaw cycles. **b** Volunteer stool sample exposed to multiple freeze–thaw cycles. **c** Human CCD time-series sample (Sample ID: 66953). **d** Human CCD time-series sample (Sample ID: LS.7.3.16). Fragment sizes (bp) are indicated in blue above each trace. **e** Histogram showing the distribution of read lengths across human fecal samples without size-selection and amplification, and **f** with size-selection and amplification. Note: the y-axis represents the number of reads and is higher in **f**.

**Extended Data Fig. 3:**
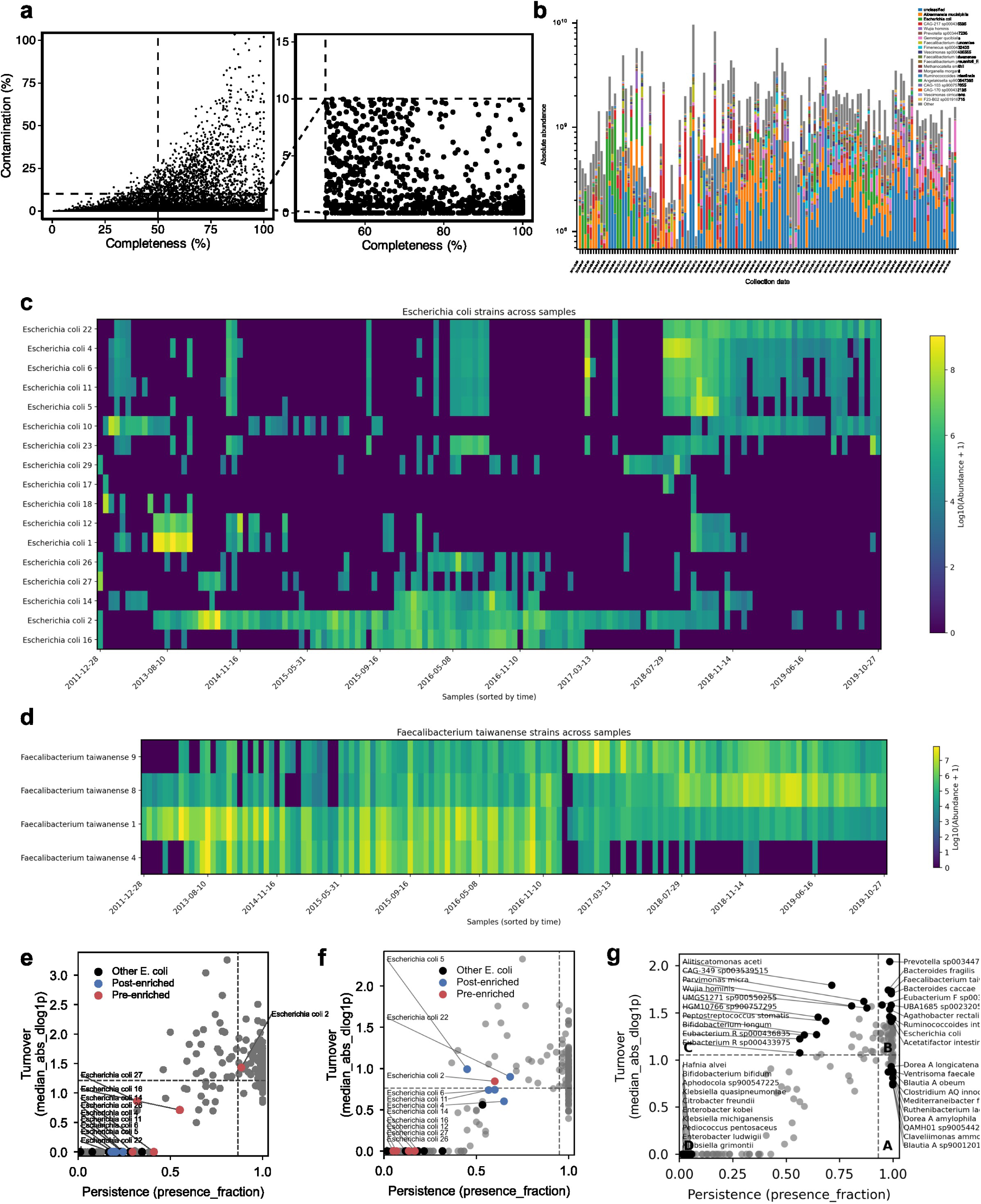
**a** Distribution of completeness and contamination across all generated LR-MAGs (left), and all LR-MAGs above 50% completeness and 10% contamination. **b** Stacked bar plot showing longitudinal log-scale absolute abundances of the dominant microbial species across all sampled collection dates. Each bar represents one stool sample, colored segments indicate individual taxa, and grey represents the aggregate “Other” category. **c** Heatmap of absolute abundances for detected *Escherichia coli* strains across samples ordered by collection date. Rows represent individual *E. coli* strains and columns represent samples, with color indicating log-transformed absolute abundance. **d** Heatmap of absolute abundances for detected *Faecalibacterium taiwanense* strains across the same ordered samples. **e** LR-MAG-level comparison of persistence and turnover for *E. coli* strains in the pre-surgery period. Persistence is defined as the fraction of samples in which a strain is detected, and turnover is summarized as median absolute log-change between observations; points are colored by whether strains are enriched before or after surgery. **f** Post-surgery view of the *E. coli* persistence–turnover space, emphasizing that several low-persistence *E. coli* strains show episodic detection, while the post-enriched subset shows broader persistence across the time series. **g** Species-level persistence–turnover analysis across the broader microbial community. Dashed thresholds separate low-persistence/low-turnover taxa from highly persistent or highly dynamic taxa, defined as the median turnover or persistence values, identifying organisms with possible stable long-term colonization versus taxa with more episodic or rapidly changing abundance profiles.

**Extended Data Fig. 4:**
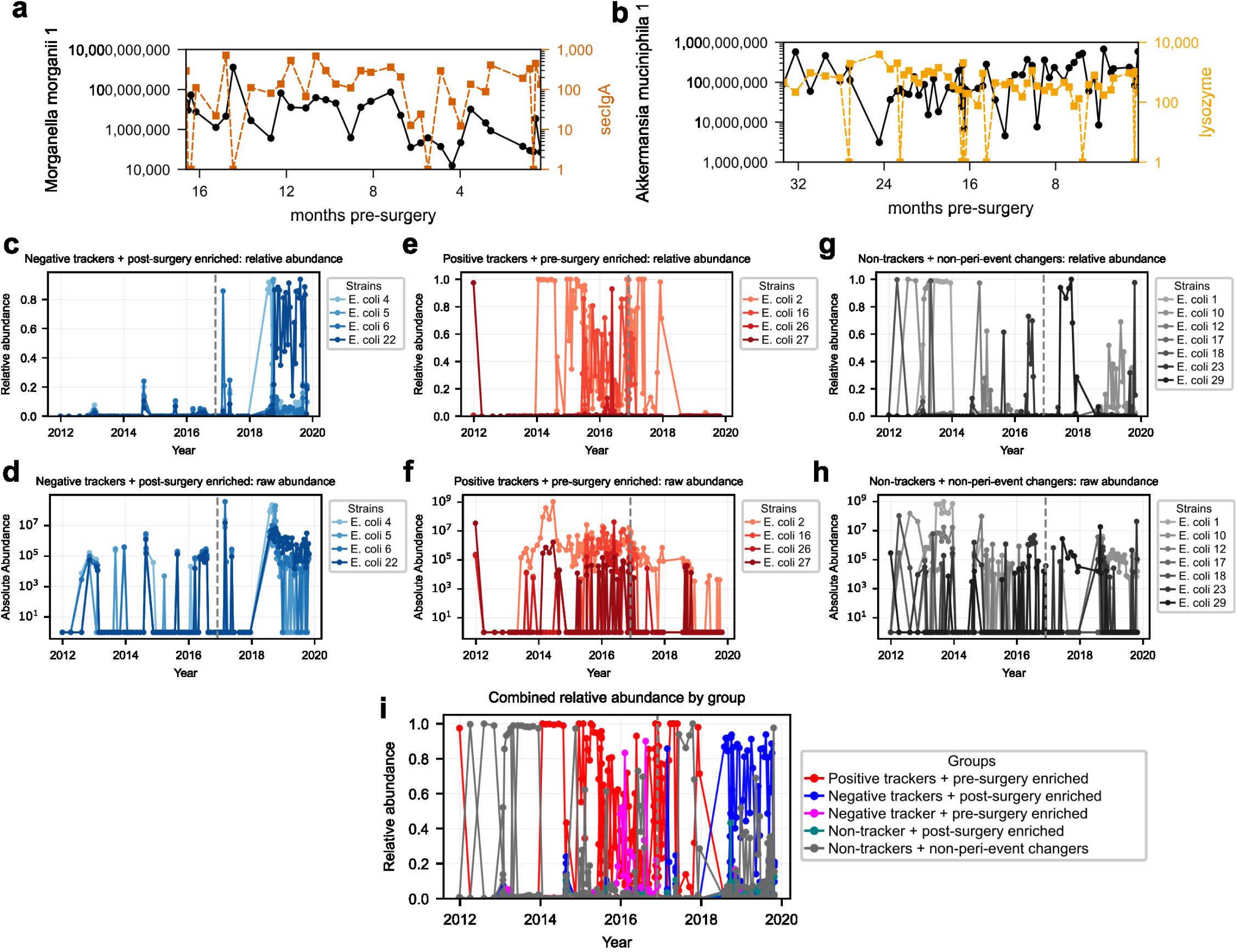
**a** Absolute abundance trajectory of *Morganella morganii* 1 during the pre-surgery period plotted against secretory IgA. The strain shows temporal covariation with the inflammatory/immune marker across the months preceding surgery. **b** Absolute abundance trajectory of *Akkermansia muciniphila* 1 during the pre-surgery period plotted against lysozyme, showing marker-associated abundance dynamics for a dominant mucus-associated gut symbiont. **c** Relative abundance trajectories of *E. coli* strains classified as negative inflammatory-marker trackers and post-surgery enriched. These strains are largely low-abundance before surgery but become dominant in the post-surgery period. **d** Absolute abundance trajectories for the same negative-tracker/post-surgery-enriched strains, showing that their post-surgery increase is also observed in absolute cell-count space rather than only as a relative-abundance effect. Importantly, the level of absolute abundances are strikingly different than what can be observed in relative abundance. **e** Relative abundance trajectories of positive inflammatory-marker trackers that are enriched before surgery. These strains dominate during the high-inflammation pre-surgery interval and decline after surgery. **f** Absolute abundance trajectories for the same positive-tracker/pre-surgery-enriched strains, showing elevated absolute abundance before surgery with reduced detection or lower abundance after the peri-surgical transition. **g** Relative abundance trajectories of *E. coli* strains classified as non-trackers and non-peri-event changers. **h** Absolute abundance trajectories for the same non-tracker/non-peri-event-changing strains. **i** Combined relative abundance trajectories of all *E. coli* strain groups, summarizing the temporal separation between pre-surgery positive trackers, post-surgery negative trackers, and strains without consistent marker-tracking or peri-event enrichment. The dashed vertical line denotes the surgery-associated transition point.

**Extended Data Fig. 5:**
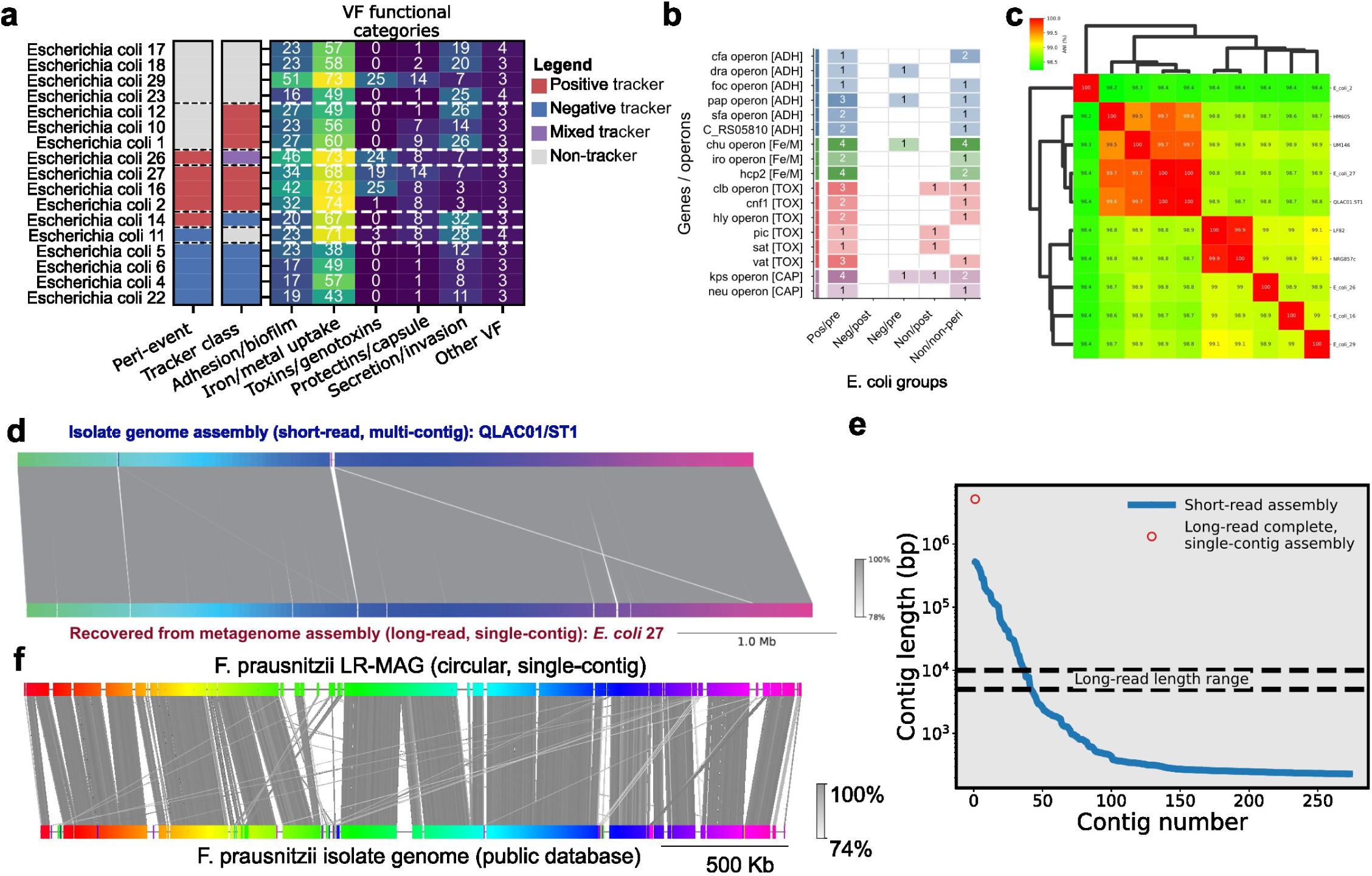
**a** Virulence-factor functional profiles of *E. coli* strains grouped by peri-event enrichment and inflammatory-marker tracking class. Rows represent individual *E. coli* strains, side annotations indicate peri-event and tracker classifications, and heatmap values summarize the number of detected virulence features in each functional category. **b** Distribution of selected virulence genes and operons across *E. coli* strain groups. Features are grouped by functional class, including adhesion, iron/metal uptake, toxins/genotoxins, and protectins/capsule, highlighting group-specific enrichment of pathogenicity-associated loci. **c** Pairwise average nucleotide identity clustering of selected *E. coli* strains and reference genomes, showing that LRQ-resolved strains span closely related but genetically distinguishable *E. coli* lineages, including IBD-isolated pathogenic strains. **d** Whole-genome alignment between a complete, single-contig long-read genome of *E. coli* 27 recovered from metagenome assembly and a multi-contig, short-read assembled genome for QLAC01/ST1, isolated from the same patient, shows near-complete colinearity and high nucleotide identity. **e** Comparison of the size distribution of the contigs comprising the short-read isolate genome assembly of QLAC01/ST1 and the single-contig size of the long-read metagenome assembled LR-MAg of *E. coli* 27. **f** Whole-genome alignment between a complete, single-contig long-read genome of *F. prausnitzii* recovered from metagenome assembly and a long-read assembled genome of *F. prausnitzii* from public databases (GCF_019967995.1).

**Extended Data Fig. 6:**
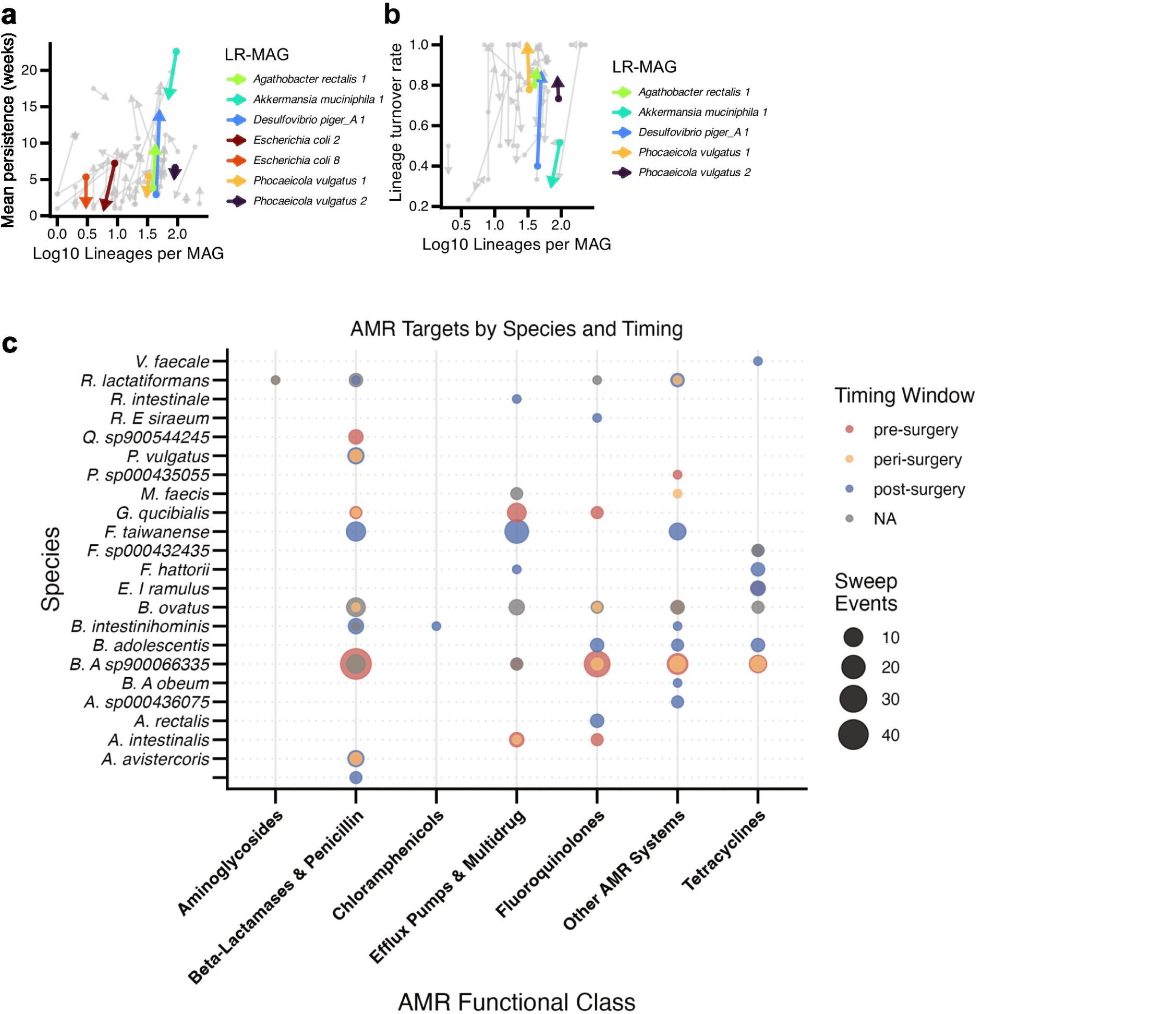
**a** MAG-level lineage persistence before and after surgery. Each point represents a MAG summarized by the log-transformed number of distinct detected lineages and the mean persistence of those lineages in weeks. Arrows connect the same MAG between pre-surgery and post-surgery phases. Selected species, including *Akkermansia muciniphila*, *Agathobacter rectalis*, *Dysosmobacter piger*, *Escherichia coli*, and *Phocaeicola vulgatus*, exhibit different peri-event abundance and marker tracking, and are colored to emphasize species-specific changes. **b** MAG-level lineage turnover before and after surgery. Each point represents a MAG summarized by the log-transformed number of distinct detected lineages and the lineage turnover rate, defined as the fraction of observed weeks in which the dominant lineage was replaced by another lineage. Arrows connect pre-surgery and post-surgery values for the same MAG, showing species-specific shifts in dominant-lineage replacement. **c** AMR-associated sweep events summarized by microbial species, AMR functional class, and timing window. The x-axis indicates AMR target class, the y-axis indicates species, point size reflects the number of sweep events, and point color denotes whether sweeps occurred pre-surgery, peri-surgery, post-surgery, or outside a defined timing window. This analysis highlights that AMR-linked sweeps are distributed across multiple taxa and resistance classes, with some species showing timing-specific enrichment of resistance-associated evolutionary events.

**Extended Data Fig. 7:**
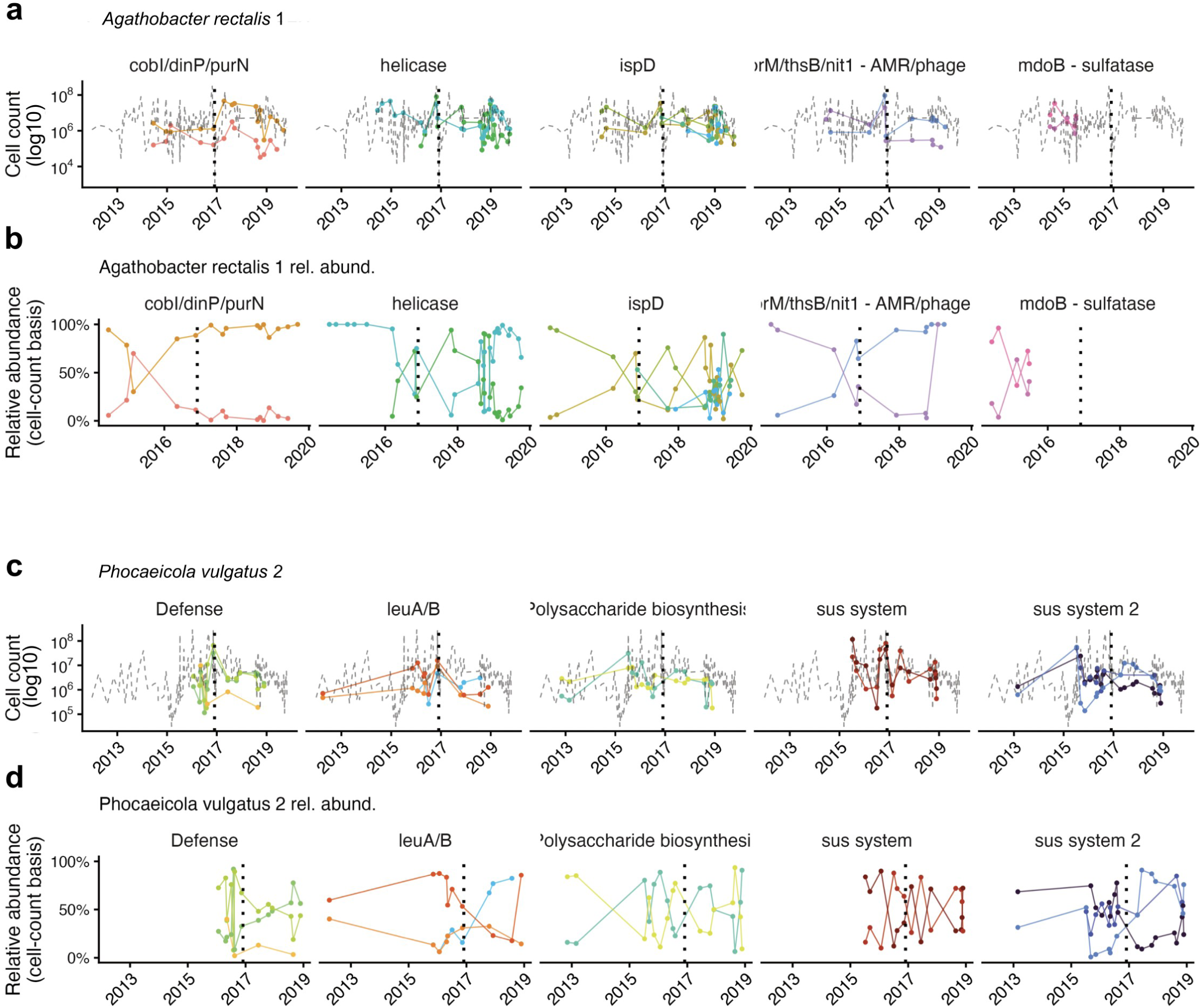
Lineage dynamics of LR-MAGs. **a** Dynamics tracked across four functionally annotated loci showing absolute haplotype cell abundances (log₁₀ CFU) over time and **b** relative haplotype abundance for LR-MAG *A. rectalis* 1. **c** Dynamics tracked across four functionally annotated loci showing absolute haplotype cell abundances (log₁₀ CFU) over time and **d** relative haplotype abundance for LR-MAG *P. vulgatus* 2.

## SUPPLEMENTARY MATERIALS

Extended Data Figures 1-7, and Extended Data Tables 1-2

## ACKNOWLEDGEMENTS

We would like to thank the Stem Cell Genomics Core at the Sanford Stem Cell Institute for providing sequencing services. We thank Dr. Trevor Biddle, Genomics Core Scientist, for his helpful discussions during the course of this work. We would also like to thank Pacific Biosciences for technical advice on PacBio SMRTbell preparation protocols and data demultiplexing. This publication includes data generated at the UC San Diego IGM Genomics Center utilizing an Illumina NovaSeq 6000 that was purchased with funding from a National Institutes of Health SIG grant (#S10 OD026929). A number of panels were created in BioRender. Din, O. (2025) https://BioRender.com/.

## CODE AVAILABILITY

The Strainphase pipeline for strain-level longitudinal metagenomics is available at https://github.com/rolesucsd/strainphase. Analyses performed using publicly available software are specified with versions throughout the Methods.

## DATA AVAILABILITY

*Raw sequencing data:* Long-read metagenomic data are available on Qiita, Study ID#10283, Prep ID’s 19140 and 19350.

## AUTHOR CONTRIBUTIONS

All authors contributed to this effort and reviewed and approved the manuscript.

